# Genomic evolution of non-small cell lung cancer patient-derived xenograft models

**DOI:** 10.1101/2023.01.06.521078

**Authors:** Robert E. Hynds, Ariana Huebner, David R. Pearce, Ayse U. Akarca, David A. Moore, Sophia Ward, Kate H.C. Gowers, Takahiro Karasaki, Mark S. Hill, Maise Al Bakir, Gareth A. Wilson, Oriol Pich, Monica Sivakumar, Assma Ben Aissa, Eva Grönroos, Deepak Chandrasekharan, Krishna K. Kolluri, Rebecca Towns, Kaiwen Wang, Daniel E. Cook, Leticia Bosshard-Carter, Cristina Naceur-Lombardelli, Andrew J. Rowan, Selvaraju Veeriah, Kevin Litchfield, Sergio A. Quezada, Sam M. Janes, Mariam Jamal-Hanjani, Teresa Marafioti, TRACERx Consortium, Nicholas McGranahan, Charles Swanton

## Abstract

Patient-derived xenograft (PDX) models of cancer, developed through injection of patient tumour cells into immunocompromised mice, have been widely adopted in preclinical studies, as well as in precision oncology approaches. However, the extent to which PDX models represent the underlying genetic diversity of a patient’s tumour and the extent of on-going genomic evolution in PDX models are incompletely understood, particularly in the context of heterogeneous cancers such as non-small cell lung cancer (NSCLC). To investigate the depiction of intratumour heterogeneity by PDX models, we derived 47 new subcutaneous multi-region PDX models from 22 patients with primary NSCLC enrolled in the clinical longitudinal cohort study TRACERx. By analysing whole exome sequencing data from primary tumours and PDX models, we find that PDX establishment creates a genomic bottleneck, with 76% of PDX models being derived from a single primary tumour subclone. Despite this, multiple primary tumour subclones were capable of PDX establishment in regional PDX models, indicating that PDX libraries derived from multiple tumour regions can capture intratumour heterogeneity. Acquisition of somatic mutations continued during PDX model expansion, and was associated with APOBEC- or mismatch repair deficiency-induced mutational signatures in a subset of models. Overall, while NSCLC PDX models retain truncal genomic alterations, the absence of subclonal heterogeneity representative of the primary tumour is a major limitation. Our results emphasise the importance of characterising and monitoring intratumour heterogeneity in the context of pre-clinical cancer studies.

## INTRODUCTION

In patient-derived xenograft (PDX) models, human tumours are propagated by transplantation into immunocompromised mice^1^. PDX models have become important models in cancer biology as they are thought to mimic tumour biology more closely than traditional cell lines as a consequence of their *in vivo* cell-cell and/or cell-matrix interactions, 3D architecture and relatively recent derivation^2^. Many reports have suggested that the responses of PDX models to drug treatment are concordant with those observed in patients, either at the level of histological subtypes or at the level of individuals. The former has led to the use of PDX models in pre-clinical drug trials prior to patient investigations^3^, while the latter has provided a personalised medicine approach, in which PDX models are used as ‘avatars’ for individual patient responses to therapy in ‘co-clinical’ trials^4,5^.

For pre-clinical oncology applications, the fidelity of PDX models is of major importance. Across cancer types, including non-small cell lung cancer (NSCLC)^6^, PDX models bear histological similarity to the tumours from which they were derived. However, recent high-resolution analyses of breast cancer PDX models suggest that PDX models, like patient tumours, can comprise multiple genetically defined subclones^7^ and that these undergo dynamic changes in their relative abundance during PDX engraftment and expansion^8^. Moreover, analysis of PDX model copy number profiles has cast doubt upon their representation of tumour molecular heterogeneity, specifically with regard to genomic evolution within the mouse^9,10^. While some of these differences may be attributable to technical issues surrounding the estimation of copy number profiles from RNA sequencing data, disagreement about the extent and importance of PDX copy number divergence remains when considering DNA sequencing data^10,11^. While some studies have included examples of matched patient-PDX pairs or the derivation of multiple PDX models from the same tumour, the genomic evolution during PDX model establishment and propagation has not been systematically assessed, the role of spatial sampling is unexplored in this context and no study to date has been performed in the context of multi-region patient sequencing data to formally establish how well PDX models represent the complex subclonal nature of primary tumours and their metastases.

Lung TRACERx is a prospective cohort study that aims to characterise the evolutionary dynamics of NSCLC through a multi-region deep whole-exome sequencing (WES) approach^12^. Here, we derive PDX models from multiple regions of primary NSCLC from patients enrolled in lung TRACERx to determine the histological and genetic fidelity of the PDX approach. By subjecting these PDX models to deep WES and comparing this dataset to multi-region WES data from matched primary tumours, we investigate key unresolved issues in the use of PDX models, including the extent of genomic bottlenecking upon engraftment, the reproducibility of PDX derivation across spatially distinct replicate samples and the emergence of *de novo* genetic alterations in PDX models over time during propagation in mice.

## RESULTS

### PDX model establishment is variable in multiply-sampled NSCLC tumours

Primary NSCLCs from patients enrolled in the lung TRACERx study undergo multi-region WES using a defined sampling protocol^12^. To characterise tumour evolution during PDX model engraftment and propagation, we obtained matched region-specific tumour material and created patient-matched PDX models from a representative patient subset (Figure 1A; Supplementary Figure 1). 145 specimens from 44 patients undergoing surgical resection of their primary NSCLC were injected subcutaneously in NOD/SCID/IL2Rg^-/-^ (NSG) mice, generating 63 xenografts from a cohort representing most NSCLC clinical and molecular subtypes (Figure 1B). Either fresh or cryopreserved tumour material was used to initiate xenografts, with no observed effect of prior cryopreservation on engraftment efficiency (p = 0.686, Chi-square test; Supplementary Figure 2A). Quality control for the presence of human lymphocytic tumours^13,14^ revealed that 16 xenografts were human CD45-expressing lymphoproliferations rather than keratin-expressing NSCLCs (Supplementary Figure 2B). One case (CRUK0885 Region R3) lacked expression of either keratin or hCD45 but was deemed NSCLC as the immunophenotype and tumour morphology was consistent with the diagnosed primary tumour subtype of carcinosarcoma. hCD45-expressing cells were absent from first generation NSCLC PDX models in all cases except CRUK0816 R2, where the number of CD45+ cells declined over passages (Supplementary Figure 2C). Thus, our NSCLC PDX cohort consisted of 47 xenografts from 22 patients with a successful engraftment rate of 50% at the tumour level and 32.4% at the region level (Figure 1B, lower panel). A bootstrapping approach suggested that single region sampling within our cohort would have resulted in PDX models for a median of 14 patients (Supplementary Figure 2D). Multiple, spatially distinct NSCLC PDX models were established for 9 patients (median = 4 regional PDX models per patient; Figure 1B). Mice with no apparent xenograft were terminated after a median of 306 days (range 37-402 days; Supplementary Figure 2E). Each region-specific PDX model was propagated by transfer of xenograft fragments to naïve hosts, maintaining the models independently, exclusively *in vivo* and generating a large biobank of cryopreserved PDX tissue. PDX models could be re-established following cryopreservation (Supplementary Figure 2F). Initial P0 PDX models took a median of 91 days before tumour harvest (range 37 - 440 days; Figure 1C), with no effect of primary tissue cryopreservation observed. In subsequent passages, PDX growth was more rapid, with a median time to harvest of 54 days across passages P1-P3 (median values for P1, P2 and P3 were 53.5, 56.5 and 51.0, respectively; Figure 1C).

**Figure 1:**
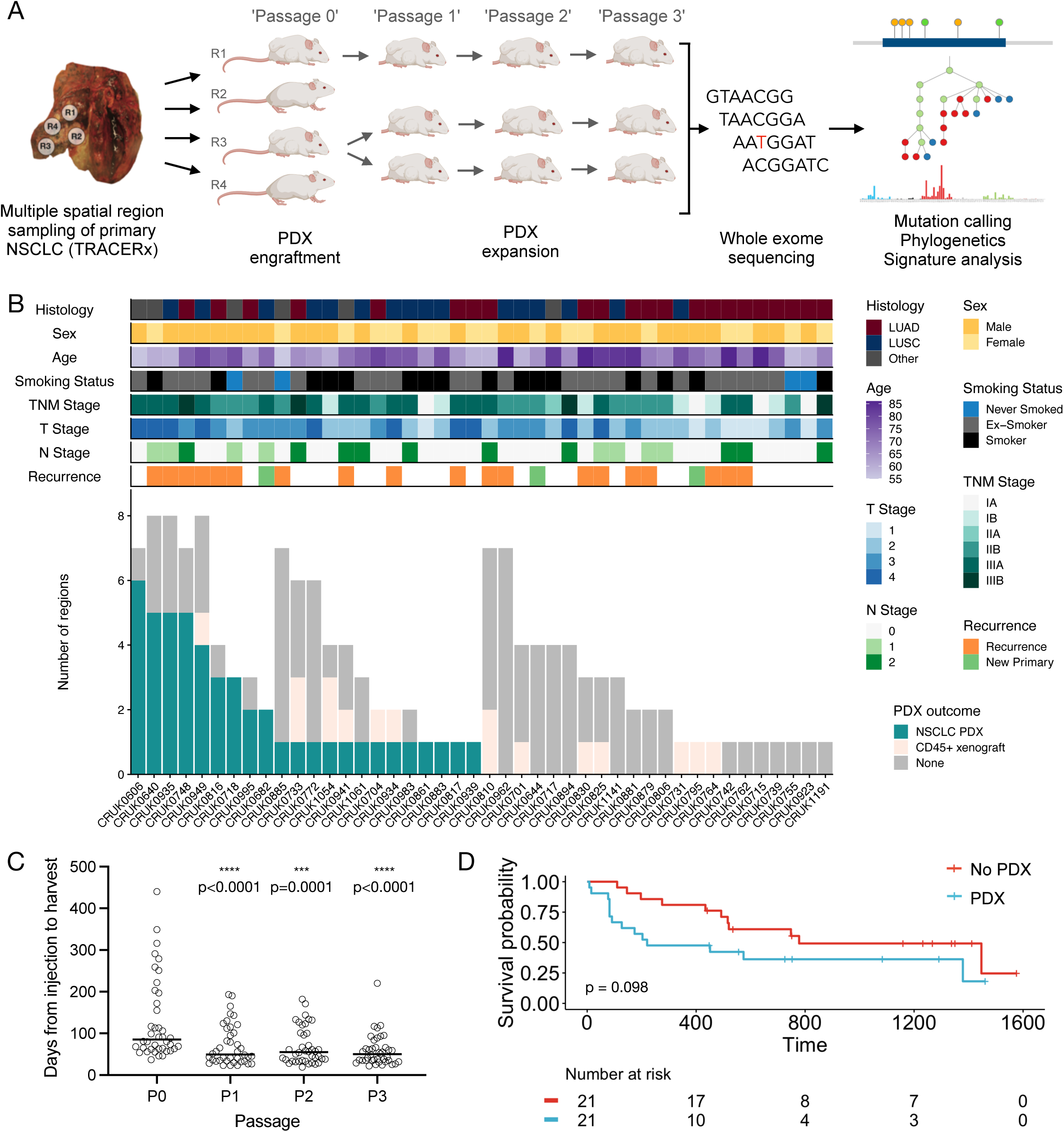
Lung TRACERx patient-derived xenograft (PDX) cohort overview. A) Schematic of study protocol to derive and expand PDX models within the lung TRACERx study. B) Outcomes of regional xenografts with patient characteristics. C) Time from tumour injection to PDX harvest by passage number. Only PDX models for which complete P0-P3 data were available are shown. Bar shows median time for all models. *P* values obtained using a Friedman test with Dunn’s test for multiple comparisons. **** p < 0.001 compared to P0. D) The proportion of patients who were disease-free over a 1600 day period following tumour resection is shown grouped by the generation (PDX) or not (no PDX) of at least one regional PDX model for each patient.

We observed a trend towards shorter disease-free survival in patients for whom at least one PDX model was established (Figure 1D). Although there was a trend towards higher engraftment from tumours with a higher T stage (Chi-square test, p=0.077), univariate analysis of clinical characteristics showed no significant differences in sex, smoking pack years, pleural invasion, vascular invasion, N stage or TNM stage, but showed that patient age and lesion size were significantly associated with PDX engraftment (Supplementary Figure 3A-3I). Seven of 24 (29.2%) lung adenocarcinoma (LUAD) tumours engrafted compared to ten of 16 (62.5%) lung squamous cell carcinoma (LUSC) tumours (p = 0.053, two-tailed Fisher’s exact test; Supplementary Figure 3J), consistent with literature reports of greater engraftment rates for LUSC histology NSCLCs^15–22^. However, this patient-level analysis is complicated by our multiple sampling of tumours; when considering engraftment by tumour region, 14/52 LUAD (24.5%) and 18/60 LUSC (30.0%) regions formed PDX models (p = 0.835, two-tailed Fisher’s exact test; Supplementary Figure 3J).

Leveraging primary tumour sequencing data of the 44 patient PDX cohort, we found no differences in the overall number of mutations, the proportion of truncal and subclonal mutations, the proportion of truncal and subclonal copy number alterations, or mutational signatures in primary tumours which yielded at least one PDX model compared to those which did not (Figure 2; Supplementary Figure 4A-4D). Assessing the presence or absence of specific driver mutations revealed that *TP53* mutations were enriched in tumours that gave rise to PDX models compared to those that did not (Fisher’s exact test, p=0.015; Figure 2; Supplementary Figure 4E). The tumour purity of engrafted regions was higher than for non-engrafted regions in patient tumour exome sequencing data (Supplementary Figure 4F) and correspondingly, T cell infiltration of the primary tumour regions was lower for engrafted regions as estimated using the T cell ExTRECT tool^23^ (Supplementary Figure 4G), supporting the view that tumour sampling (likely the absolute viable tumour cell number injected) is a major factor determining engraftment success^24^. Additionally, when considering copy number metrics, we find that tumours that gave rise to PDX models had a higher fraction of the genome subject to loss of heterozygosity (Supplementary Figure 4H), although there was no difference in weighted genome instability index (Supplementary Figure 4I).

**Figure 2:**
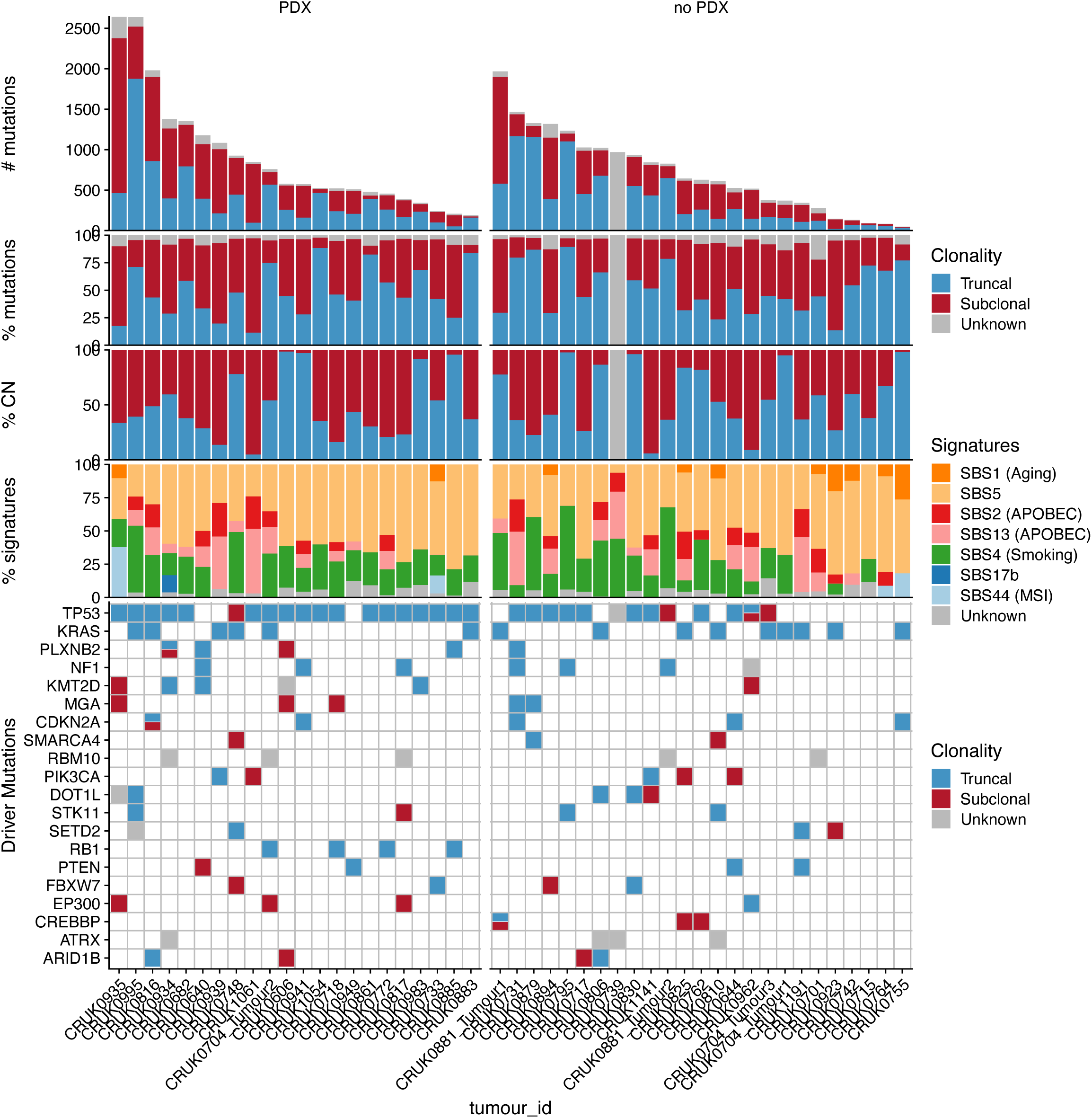
Genomic characteristics of primary tumours. Patient primary tumours are split based on whether a PDX model was engrafted from any tumour region (PDX) or not (no PDX). Within each category, tumours are ordered according to their total mutation burden. Top panel: total number of coding and non-coding mutations including SNVs, dinucleotide and indel alterations. Bars are coloured by the clonality status of alterations. Second panel: proportion of truncal and subclonal mutations. Third panel: proportion of truncal and subclonal copy number alterations. Fourth panel: proportion of mutational signatures as estimated across all mutations. Bottom panel: driver alterations on a per tumour basis. The mutations shown are the 20 most frequently mutated genes in this patient cohort. Mutations are coloured by the clonal status of alterations.

For a majority of region-specific PDX models, there were strong histological similarities between the patient sample and both the early and late PDX tissues (Supplementary Figure 5A), suggesting stability during serial engraftment. However, in some cases, we noted histological variation. Some models diverged immediately; for example, CRUK0949 R1 and R3 showed more widespread clear cell differentiation than was present in the corresponding patient samples for those regions, and CRUK0816 R2 and R5 PDX models presented more epithelioid differentiation than the parent tumour (Supplementary Figure 5B). Other models varied between early and late passage with the P0 xenograft more closely resembling the patient region than the P3 xenograft; for example, CRUK0941 R2 PDX model showed prominent rhabdoid differentiation in hematoxylin and eosin stained sections at the later time point that had not been present in either the patient or early passage samples (Supplementary Figure 5C), though this is consistent with the cytological pleomorphism seen in this poorly differentiated pleomorphic carcinoma. In different CRUK0606 regional PDX models, variation between either tumour and P0 PDX models, or P0 and P3 PDX models were observed. Glandular features were a minor component of the patient’s regional tissue but became more prominent in PDX models, either in both early and late passage models (R5, R8) or in the late passage model only (R1, R6; Supplementary Figure 5C).

### Genomic bottlenecks associated with engraftment lead to monoclonality of PDX models

We performed WES on PDX models once at their first establishment in mice (“passage zero”, P0; median of 91 days after initial injection) and again at passage three (a median of 279 total days since P0 initial injection; Figure 1C), for comparison to primary regions in the TRACERx study. WES data were filtered to remove contaminating mouse reads using the bamcmp tool^25,26^ and, initially, the mouse GRCm38 (mm10; C57BL/6J strain) reference genome. This resulted in mouse strain-dependent erroneous inclusions in mutation calling, which could not be confirmed by Sanger sequencing, leading us to develop an NSG-adapted reference genome that improved the accuracy of mouse read removal.

In the knowledge that primary tumour regions generally consist of multiple subclones, we inferred the subclonal composition of P0 PDX tumours relative to their primary tumour region of origin. If a single subclone was shared between the primary tumour region and the P0 PDX, we defined this as monoclonal PDX engraftment; if multiple subclones were found, we defined this as polyclonal PDX engraftment (Figure 3A). In PDX models for which the clonality status of the region of origin was known, 21 of 26 models from heterogenous primary tumour regions were monoclonal, suggesting a major bottleneck during PDX engraftment (Figure 3B; Supplementary Figure 6A). A further five monoclonal PDX models arose from homogenous primary tumour regions, while 10 PDX models from heterogenous showed polyclonal engraftment (Figure 3B; Supplementary Figure 6A).

**Figure 3:**
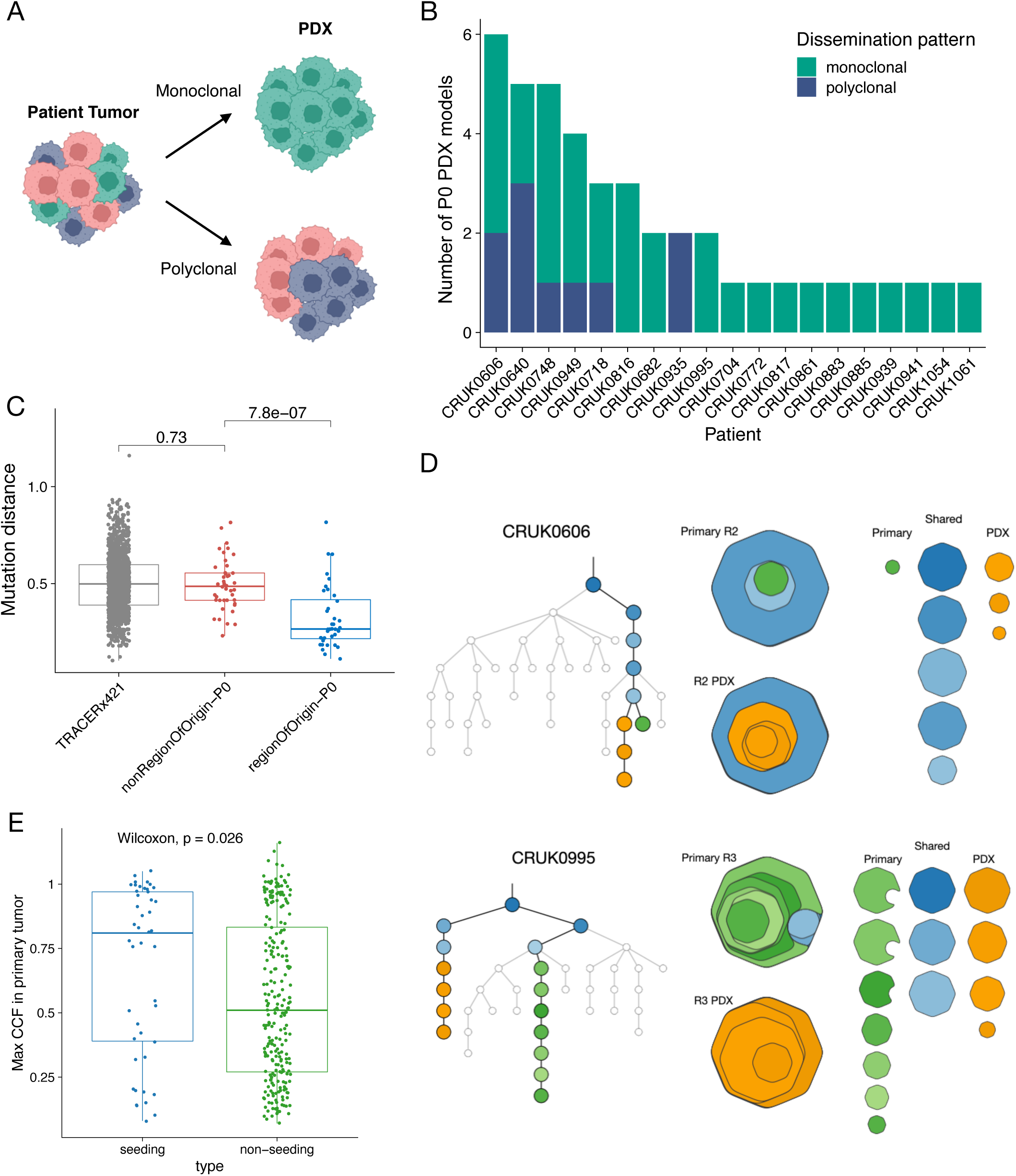
Genomic comparison of primary tumour region-early passage PDX pairs. A) Schematic representation of dissemination patterns. B) Overview of dissemination patterns (monoclonal, green; polyclonal, blue) for each early passage (P0) PDX sample for which WES data were available. Data are ordered by total number of PDX samples. C) Mutational distance between regions within each primary tumour in the lung TRACERx421 cohort, P0 PDX models and other regions of their primary tumour and P0 PDX models and their respective region of origin. *P* values obtained by two-sided Wilcoxon rank sum test. D) Examples of comparisons of P0 PDX models and their region of origin. The upper panel is the CRUK0606 R2 PDX model, the lower panel is the CRUK0995 R3 PDX model. These were selected as the greatest and lowest mutational distances within the cohort, respectively. E) Maximum cancer cell fraction (CCF) of the primary tumour across all regions for PDX engrafting clusters and non-engrafting clusters. *P* values obtained by two-sided Wilcoxon rank sum test.

To explore how representative multi-region NSCLC PDX models are of intratumoural heterogeneity, we calculated a mutational distance score (see Methods) for each PDX model compared to its region of origin and to all other spatially distinct regions from the same tumour. We also calculated the mutational distance between regions for each primary tumour in TRACERx421 data. PDX models were significantly more similar to their region of origin than to spatially distinct tumour regions within the same tumour with the distance to non-region of origin comparable to that between primary tumour regions (Figure 3C; median non-region of origin-P0 = 0.486 [IQR 0.414-0.555] versus median region of origin-P0 = 0.266 [IQR 0.216-0.417]; p = 7.8×10^−7^, two-sided Wilcoxon rank sum test). However, we observed notable variability in the extent of similarity to the region of origin in different cases. At one extreme, the CRUK0606 R2 P0 PDX model was highly similar to its region of origin, with the lowest mutational distance within the cohort; the majority of clusters were shared, with only a small number of mutations differing between the two (Figure 3D, upper panel). Conversely, the mutations shared between the CRUK0995 R3 primary tumour region and the matched P0 PDX were low frequency within the primary tumour, and both the primary tumour region and P0 PDX contained many additional mutations that were not shared within WES data (Figure 3D, lower panel). This highlights that PDX models are more representative of their region of origin than other regions from the primary tumour and validates the approach of multi-region PDX generation. Consistently, we observed that mutational distance was correlated with a copy number distance metric (see Methods; Supplementary Figure 6B; Pearson’s correlation, *R* = 0.68, p = 6.3e-06). Next, we linked the mutational distance to PDX engraftment clonality and found that, by virtue of harbouring more clones from their primary regions of origin, PDX models subject to polyclonal engraftment exhibited a lower mutation distance to their respective region of origin than models that exhibited monoclonal engraftment (Supplementary Figure 6C; median monoclonal = 0.29 [IQR 0.23-0.45] versus median polyclonal = 0.238 [IQR 0.182-0.262] ; p = 0.053, two-sided Wilcoxon rank sum test). We assessed the maximum cancer cell fraction (CCF) of clusters across all primary tumour regions to determine the size of the clones that engrafted in PDX models, finding that engrafted clones had higher CCF values than non-engrafted clones (Figure 3E; median seeding = 0.76 [IQR 0.322-0.905] versus median non-seeding = 0.51 [IQR 0.27-0.832]; p = 0.026, two-sided Wilcoxon rank sum test).

In order to classify PDX models whose bottleneck event upon engraftment was characterised by predominantly mutation versus copy number events, both distances were z-transformed for normalisation. PDX models from the upper and lower quartiles of the difference between mutation and copy number distances were classified as higher mutation or copy number diversity, respectively (Supplementary Figure 6D). One regional PDX model from CRUK0748 (R1) was found in the higher mutational diversity category, while three other regions (including R6) had higher copy number diversity. Consistent with this, more substantial copy number differences, including mirrored subclonal allelic imbalance (MSAI) on chromosome 3q, were observed between the CRUK0748 R6 primary region and P0 PDX model (Supplementary Figure 6E).

Overall, these data suggest a model in which PDX engraftment induces a genomic bottleneck that commonly results in a single tumour subclone engrafting in PDX models. By inference, it is clear that PDX models do not represent the full subclonal diversity of the primary tumour. Therefore, since models are similar to their regions of origin, collections of PDX models might go further towards fully recapitulating intratumour heterogeneity for individual patients through attempts at multi-region PDX engraftment (e.g. CRUK0606; Figure 4).

**Figure 4:**
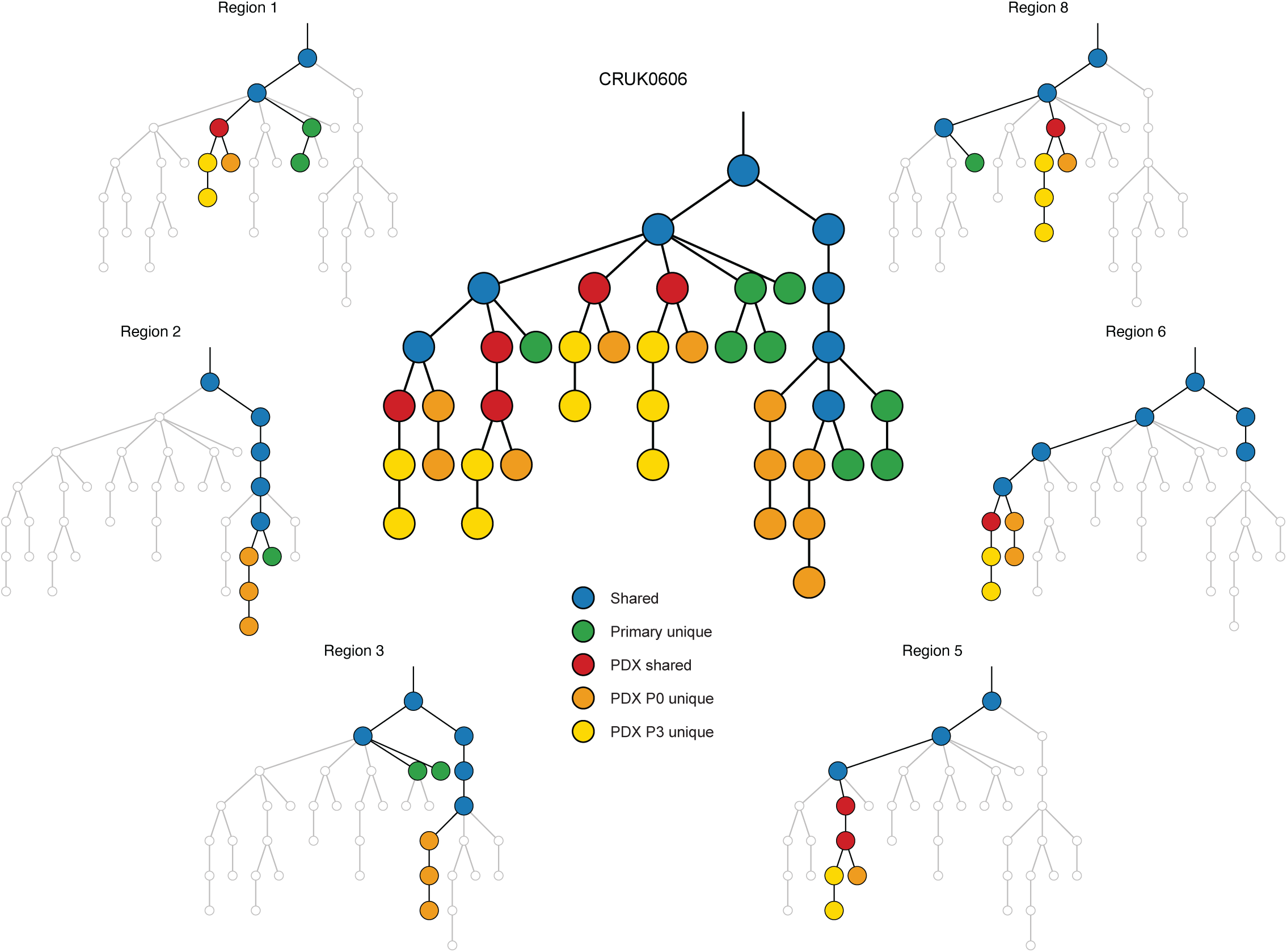
Phylogenetic tree for CRUK0606 with trees for individual PDX models.

### Propagation of PDX models involves on-going genome evolution

The phylogenetic analysis of PDX models also revealed multiple clusters of PDX unique mutations (Figure 4), suggestive of on-going evolution distinct from the primary tumour region from which the PDX model was derived. Comparison of ‘late’ passage P3 models and initial P0 models revealed that nine of ten polyclonal P0 models were monoclonal with respect to the primary tumour at P3 (Figure 5A), indicating that PDX models often do not retain subclonal complexity reflective of the primary tumour region. Consistent with this, comparing the mutational distance between the region of origin and P0 PDX pairs with P0-P3 PDX pairs showed that P3 PDX models were more similar to P0 than P0 were to the region of origin (Figure 5B; median region of origin-P0 = 0.265 [IQR 0.216-0.361] versus median P0-P3 = 0.192 [IQR 0.154-0.234]; p = 1e-04, two-sided Wilcoxon rank sum test), likely due to the substantial initial genetic bottleneck (Figure 3). PDX models where sequential WES was performed supported the notion that initial engraftment represented a strong bottleneck in terms of both mutational (Supplementary Figure 7A) and copy number (Supplementary Figure 7B) diversity, but PDX models were stable once established.

**Figure 5:**
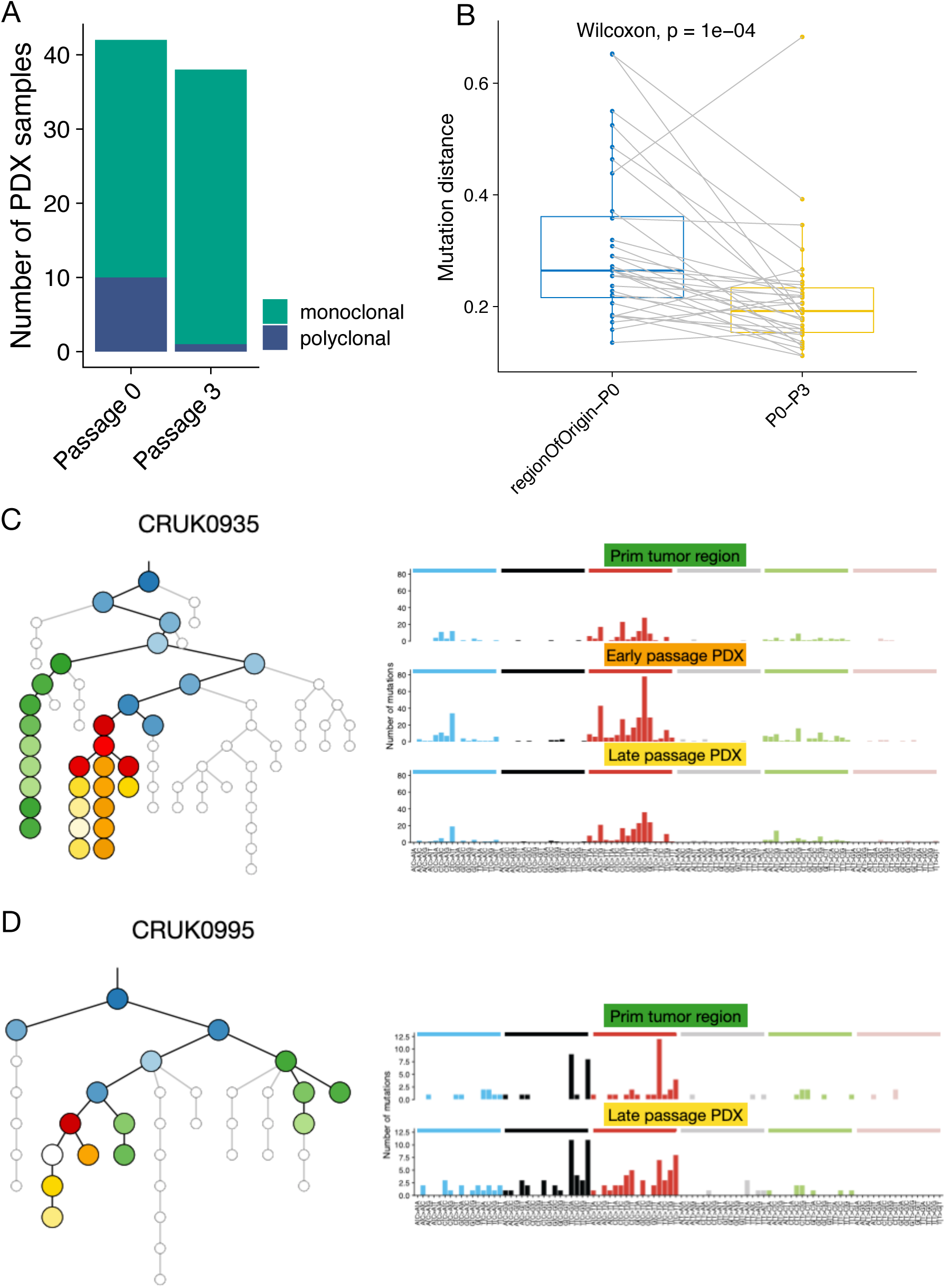
On-going evolution in NSCLC PDX models. A) Overview of dissemination patterns relative to the primary tumour of early (P0) and late (P3) PDX models. B) Comparison of mutational distance between P0 PDX models and the region of origin, and P3 PDX models with the corresponding P0 PDX model. *P* values obtained by two-sided Wilcoxon signed rank test. C) Matched primary tumour region, early (P0) PDX and late (P3) PDX mutational signature analysis for CRUK0935 R5. On the phylogenetic tree (left), shared clusters between the primary tumour and PDX model are coloured blue, those found in the primary tumour region only are coloured green, shared mutation clusters between early and late PDX models are coloured red, while those found in the P0 or P3 PDX models only are coloured orange or yellow-white, respectively, with different gradient colours denoting different mutational clusters. Analysis of mutational context revealed a signature reminiscent of mismatch repair deficiency in primary tumour region-unique, P0 PDX-unique and P3 PDX-unique mutations. D) Matched primary tumour region and P3 PDX mutational signature analysis for CRUK0995 R1. On the phylogenetic tree (left), shared clusters between the primary tumour and PDX model are coloured blue, those found in the primary tumour only are coloured green, shared mutations between early and late PDX models are coloured red, while those found in the P0 or P3 PDX models only are coloured orange or yellow, respectively. Analysis of mutational context revealed a signature consistent with APOBEC mutagenesis in primary tumour region-unique and P3 PDX-unique mutations. There were insufficient P0 PDX-unique mutations to perform this analysis at that time point.

Some models had sufficient numbers of unique mutations to perform mutational signature analysis for both the primary tumour region of origin and PDX models. The CRUK0935 primary tumour regions were mismatch repair deficient and both R5 P0 PDX- and R5 P3 PDX-unique mutations showed evidence of an on-going MMR signature (Figure 5C). Similarly, we found that CRUK0995 R1 had evidence of APOBEC signature mutations in both the primary tumour and the matched late passage PDX model, indicative of APOBEC-induced mutagenesis during PDX expansion (Figure 5D).

## DISCUSSION

Here we have investigated the genomic evolution of NSCLC during subcutaneous engraftment and propagation in immunocompromised mice. Although a recent pan-cancer WES study found ∼10% discordance in driver mutations in patient-PDX pairs, indicative of clonal evolution during engraftment^27^, previous studies based on gene expression profiling, SNP array, panel sequencing and/or whole-genome sequencing have demonstrated widespread conservation of the genomic landscape in PDX models from a range of cancer types^6,8,28^. However, these studies have been limited in their ability to detect subclonal events by a lack of patient tumour multi-region sampling. There have also been conflicting reports about the extent of on-going genomic evolution within PDX models, with authors concluding that genetic drift in PDX models is either minimal^3,11,29^ or substantial^9,10^. To better address these issues, we prospectively developed a new PDX collection within the context of a NSCLC patient cohort for whom detailed annotation including multi-region whole exome sequencing was available for comparison.

Quality control to ensure model and data validity are key components of PDX model pipelines. Our findings regarding the formation of B lymphoproliferations are mirrored in previous PDX studies in NSCLC^13^ and other cancer types^30^. These are thought to arise from EBV-transformed B cells within transplanted material whose expansion is prevented by host surveillance but enabled following transplantation in immunocompromised mice^31^. Measures to ensure authentic engraftment of the tissue of interest in xenograft studies are therefore essential, and, since murine lymphomas can also be transferred during subsequent passaging^32^, regular surveillance for CD45+ xenografts is required. For sequencing data analysis, PDX workflows typically include a step to remove contaminating mouse reads (e.g. using bamcmp^25^, Xenome^33^ or other tools^34–36^). We identify mutation calls that arise in PDX samples as a result of NSG mouse DNA contamination that are not identified by filtering using the mm10 reference genome, which is based on the C57Bl/6J strain and has a divergent SNP profile to the NSG strain. By adapting the mm10 reference genome by spiking in these SNPs, we generated an improved filtering method but ultimately our data support the need for the derivation of a complete NSG reference genome assembly for use in xenograft studies.

Few previous studies have investigated multiple PDX models per primary tumour in the context of matched patient tumour sampling, so our multi-region tumour sampling data help to reframe the interpretation of data derived from single region PDX studies. One assumption of previous PDX studies has been that the success or failure of a single tumour region represents the behaviour of the tumour overall. However, we find that distinct spatial regions of the same tumour can have divergent outcomes in PDX models, and that engraftment is correlated with tumour purity and inversely associated with T cell infiltration, consistent with a study of breast cancer PDX models^37^. Prior studies also suggest that lung squamous carcinomas more readily give rise to PDX than lung adenocarcinomas^16–19^. While we found evidence to support this at the patient level, the proportion of regions giving rise to PDX models was similar between the two histologies, suggesting that sampling biases (e.g. higher tissue availability from larger tumours) may play a role in apparent histology-dependent changes in engraftment rates.

*TP53* mutations have been associated with better engraftment of EGFR-mutant lung adenocarcinomas in PDX models^38^, and we now provide evidence that this extends more broadly within NSCLC. Since tumours with higher chromosomal instability were more likely to give rise to PDX models, we speculate that this might represent an advantage in adapting to novel environments.

Previous studies using WES or WGS have typically found the conservation of a majority of tumour mutations in PDX models but are often limited in their ability to call subclonal mutations by a low depth of coverage. Here, using a sequencing approach sufficient to confidently identify subclonal mutations, we identify major genomic bottlenecks upon establishment of NSCLC PDX models, consistent with the findings that minor tumour subclones can dominate breast cancer xenografts^8^. In our NSCLC cohort, this rendered the majority of PDX models monoclonal with respect to a polyclonal primary tumour. Such bottlenecking represents a limitation of single region PDX models, particularly in personalised therapy approaches where fully representative tumour sampling is likely to be crucial to determine the behaviour of a tumour and somatic events that drive the acquisition of drug resistance. Consistent with findings in a recent study that compared PDX samples from the same tumour biopsy with those from independent models derived from the same patient (e.g. two metastases)^27^, PDX models more closely represented the tumour region from which they were derived than more distant tumour regions, suggesting that developing multiple PDX models per patient might be useful for personalised approaches to capture intratumour heterogeneity in a PDX collection. In contrast to data from repeated transplantation of established PDX models^8^, our data suggest that multiple primary tumour subclones are capable of PDX engraftment in different engraftment attempts, giving hope that primary tumour heterogeneity can be captured within NSCLC PDX libraries.

PDX engraftment and early passaging largely resulted in PDX models that were monoclonal with respect to the patient tumour, and initially polyclonal PDX models became monoclonal over time. However, we noted that the clonal architecture of late passage PDX models could still be complex as a result of PDX-unique mutations that are not found in the primary tumour. These mutations suggested that genomic evolution was on-going in the PDX models, and we identified models that were defined by specific mutational signatures, such as mismatch repair deficiency and APOBEC mutagenesis. Such PDX models might be useful in studies aiming to reduce the mutational rates in these contexts. However, overall, the on-going accumulation of mutations over approximately 9 months of expansion in mice contributed less to the overall genomic distance of PDX models from primary tumours than did initial bottlenecking events. Nevertheless, this finding has implications for long-term modelling using PDX models and suggests the value of generating large banks of low passage PDX models and regular screening of the cohort for acquired genomic changes. It also represents an important consideration in approaches that use PDX models to derive cell lines or organoids for further study^7,39,40^.

In summary, our study tracking cancer mutations through primary NSCLC engraftment and expansion in PDX models reveals a genomic bottleneck during engraftment that often means an individual PDX model is representative of only one subclone of the primary tumour. The full representation of truncal tumour alterations in PDX models supports their use in cohort level studies and for testing therapeutics targeting truncal events. However, the underrepresentation of subclonal heterogeneity in PDX models suggests that care should be taken in extrapolating data from single region PDX models in personalised medicine approaches^41^, where the models may not be fully representative of the primary tumour. We observed on-going evolution in PDX models but models were generally stable over passage, with *de novo* events contributing less to genomic divergence than initial bottlenecking events, supporting the expansion and banking of models, although it should be noted that we have not assessed at the cumulative effect of more than three xenograft passages.

## METHODS

### Generation and maintenance of multi-region NSCLC PDX models

Ethical approval to generate patient-derived models was obtained through the Tracking Cancer Evolution through Therapy (TRACERx) clinical study (REC reference: 13/LO/1546; https://clinicaltrials.gov/ct2/show/NCT01888601). Animal studies were approved by the University College London Biological Services Ethical Review Committee and licensed under UK Home Office regulations (project license P36565407).

Tissue from patients undergoing surgical resection of NSCLCs was immediately transported on ice from the operating room to a pathology laboratory where it was dissected for diagnostic and then research purposes. Tumour samples were dissected by a consultant pathologist such that the tissue used to generate patient-derived xenograft (PDX) models was considered to be within the same tumour region as material sequenced in the TRACERx study. In cases where region-matched tissue could not be collected for PDX studies, inter-region (IR) tumour tissue was used. Tumour samples for PDX studies were transported to the laboratory in transport medium consisting of MEM alpha medium (Gibco) containing 1X penicillin/streptomycin (Gibco), 1X gentamicin (Gibco) and 1X amphotericin B (Fisher Scientific, UK). Samples were minced using a scalpel and either resuspended in 180 ul growth factor-reduced Matrigel (BD Biosciences) for fresh injection, or frozen in ice-cold foetal bovine serum plus 10% DMSO, first to -80°C in a CoolCell (Corning) before long-term storage in liquid nitrogen.

Mice were kept in individually ventilated cages under specific pathogen-free conditions and had *ad libitum* access to sterile food and water. To generate PDX tumours, male non-obese diabetic/severe combined immunodeficient (NOD/SCID/IL2Rg^-/-^; NSG) mice were anaesthetized using 2–4% isoflurane, the flank was shaved and cleaned before tumour tissue in Matrigel was injected subcutaneously using a 16G needle. Mice were observed during recovery, then monitored twice per week for tumour growth. When xenograft tumours formed, tumour measurements were taken in two dimensions using callipers and mice were euthanized before tumours reached 1.5 cm^3^ in volume. Mice without xenograft tumours were terminated after a median of 306 days (range 37-402 days). Successfully engrafted tumours were propagated through four generations of mice, with banking of histology tissue, OCT-embedded frozen tissue and xenograft DNA at each generation. Cryopreservation of living xenograft tissue was also performed at each tumour transfer as per patient tissue.

### Histopathological characterisation

Paraffin-fixed tissue sections were routinely obtained at PDX passage by fixation of tumour fragments (approx. 3×3×3 mm in size) in 4% paraformaldehyde. Samples were fixed overnight at 4°C and stored in 70% ethanol at 4°C before being processed through an ethanol gradient using an automated pipeline and embedded in paraffin. Formalin-fixed paraffin-embedded tissue sections of PDX tumours and their equivalent primary tumour region were subjected to hematoxylin and eosin (H&E) staining or immunohistochemistry with the following antibodies; anti-CD45 (Clone HI30; Dilution 1:200; Cat No 304002); anti-keratin (Clone: AE1/AE3; Dilution: 1:100; Cat No: 13160); anti-CD3 (Clone: LN10; Dilution: 1:100; Cat No: NCL-L-CD3-565); anti-CD20 (Clone L26; Dilution: 1:200; Cat No: M0755). Optimization of the antibodies was carried out on sections of human tonsil tissues. Immunostaining was performed using an automated BOND-III Autostainer (Leica Microsystems, UK) according to protocols described previously^42^.

### Genomic profiling

DNA was extracted from PDX models at each transfer using either the PureLink Genomic DNA Mini Kit (Invitrogen) or the DNA/RNA AllPrep Kit (Qiagen). For each PDX sample, exome capture was performed on 200 ng DNA using a customised version of the Agilent Human All Exome V5 Kit (Agilent) according to the manufacturer’s protocol, as previously reported ^43^. Following cluster generation, samples were 100 bp paired-end multiplex sequenced on the Illumina HiSeq 2500 and HiSeq 4000 at the Advanced Sequencing Facility at The Francis Crick Institute, London, U.K.

### Bioinformatics pipeline

#### Alignment

Initial quality control of raw paired-end reads (100bp) was performed using FastQC (0.11.8, https://www.bioinformatics.babraham.ac.uk/projects/fastqc/) and FastQ Screen (0.13.0, https://www.bioinformatics.babraham.ac.uk/projects/fastq_screen/, flags: --subset 100000; --aligner bowtie2). Subsequently, fastp (0.20.0, flags: --length_required 36; --cut_window_size 4; -- cut_mean_quality 10; --average_qual 20) was used to remove adapter sequences and quality trim reads. Trimmed reads were aligned to the hg19 genome assembly (including unknown contigs) using BWA-MEM (0.7.17)^44,45^. Alignments were performed separately for each lane of sequencing and then merged from the same patient region using Sambamba (0.7.0)^46^ and deduplicated using Picard Tools (2.21.9, http://broadinstitute.github.io/picard/). Local realignment around INDELs was performed using the Genome Analysis Toolkit (GATK, 3.8.1)^47^. Further quality control following alignment was performed using a combination of Somalier (0.2.7, https://github.com/brentp/somalier), Samtools (1.9)^48^, Picard Tools, and Conpair (0.2).

For PDX samples, the steps above were repeated twice, aligning once to the hg19 genome assembly and once to the mm10 genome assembly. Subsequently, bamcmp^25^ was used to identify contaminating mouse reads in our xenograft data. Only reads aligning solely to hg19 or better to hg19 compared to mm10 were included in subsequent downstream processing steps.

### Subsequent processing

The downstream steps of somatic mutation calling and somatic copy number alteration detections, as well as manual quality control were performed analogously to the methods described in the TRACERx 100 manuscript^43^.

### Distinguishing multiple independent tumours from a single patient

To determine whether multiple samples were genomically related, we performed a clustering step on the mutations identified in each region. Firstly, all ubiquitous mutations were determined that had a VAF greater than 1% in all regions. If more than 10 such mutations existed, the regions were deemed genomically related. Conversely, if 10 or less mutations were shared across all regions, a clustering step using the R function *hclust* was performed on the mutation VAFs across all regions. Subsequently, the resulting clustering tree was separated into two groups to determine the regions associated with two distinct tumours. This step was repeated on the two distinct tumours, respectively to yield a maximum of four distinct tumours.

### Weighted genomic instability index

The weighted genomic instability index (wGII) score was calculated as the proportion of the genome with aberrant copy number relative to the median ploidy, weighted on a per chromosome basis^49^.

### Fraction of the genome subject to loss of heterozygosity

The fraction of the genome subject to loss of heterozygosity (fLOH) score was defined as the percentage of LOH identified in the genome.

### TRACERx mutation clustering and tree building

To reconstruct tumour phylogenetic trees of each tumour from the identified somatic mutations, we developed a novel computational method to address three key challenges in phylogenetic reconstruction: (1) scaling to a high number of primary tumour and metastasis regions per patient, (2) correcting for complex evolutionary events, including mutation losses^50^, (3) removing biologically improbable clusters that either are driven by subclonal copy number or are not biologically compatible with the inferred evolutionary tree. This novel method has been extensively benchmarked and a manuscript detailing the steps as well as its application is currently in preparation. The key steps are briefly outlined below.

Firstly, mutations were clustered based on their presence/absence across regions to determine which somatic mutations likely occurred in the same tumour subclone during tumour evolution. This pre-clustering step allows the method to scale to a large number of tumour regions and improves the accuracy of identifying mutations that are present in specific samples. Mutations are defined as absent in a given region if at least 1 mutant read is observed and are grouped together when they occur in the same set of regions. Groups containing less than five mutations are not clustered further, while all other mutation groups are subsequently clustered using PyClone (v0.13.1)^51^. This clustering step is performed analogously, as described in the TRACERx 100 manuscript^43^.

Secondly, tumour phylogenetic trees were reconstructed using the identified mutation clusters. The method aims to iteratively enumerate all possible nestings of mutation clusters based on the established pigeonhole principle and the crossing rule^52^. Often a phylogenetic tree cannot be reconstructed due to the presence of erroneous clusters that are either due to artefactual mutations or errors in the called overlapping SCNAs. Therefore, the method can identify and remove these clusters to allow the reconstruction of a phylogenetic tree. Specifically, our method aims to remove clusters whose genomic location is indicative of errors in SCNAs (i.e. mutations co-localised in the genome). Additionally, in order to obtain a phylogenetic tree that meets our criteria defined above, clusters are removed such that the smallest number of mutations possible are removed from the tree, (under a principle of parsimony). This step returns the ‘default’ phylogenetic tree.

### Classifying clonality of individual clusters in tumour regions

Mutation clusters were classified as clonal, subclonal, or absent in every tumour region based on comparison with the phyloCCF estimates of the clonal cluster.

In every tumour region, the phyloCCF of mutations for each cluster of interest was compared to the clonal cluster. If no significant difference between the cluster of interest and the clonal cluster was observed (one-sided Wilcoxon test, p-value = 0.05), this cluster was defined as clonal within that region. Additionally, a lower threshold of the clonal cluster was calculated as the lower bound of the 95% confidence interval of the clonal cluster phyloCCF, up to a minimum of 0.9. If the upper bound of the 95% confidence interval phyloCCF of the cluster of interest overlapped with this lower threshold, the cluster was also defined as clonal. Conversely, if the phyloCCF estimates of the cluster of interest were significantly lower than the estimates of the clonal cluster (p-value < 0.05), and the upper bound of the 95% confidence interval of the cluster of interest was lower or equal to the lower threshold of the clonal cluster, the cluster of interest was defined as not clonal. Finally, if the mean phyloCCF of the cluster of interest was greater than 0 this cluster was defined as subclonal, otherwise the cluster was defined as absent from the tumour region.

Based on these definitions of clonality for individual clusters in all tumour regions, clonality of individual clusters could be defined within the primary tumour and across all PDX samples. Clusters that were clonal in all regions of interest (i.e. all primary regions, or PDX samples) were defined as clonal within the primary tumour or PDX samples, respectively. Clusters that were subclonal or absent from at least one region of interest were defined as subclonal, while clusters that were absent from all regions of interest were defined as absent at the tumour level.

### Analysis

#### Classifying dissemination patterns

Within each primary tumour we identified which cancer clone(s) were involved in PDX engraftment and classified the dissemination pattern as monoclonal, if only a single clone of the primary tumour was engrafted in PDX samples, or polyclonal, if multiple cancer clones were involved in engraftment.

Specifically, for each individual PDX sample, if all mutation clusters shared between the primary tumour and the sample were found to be clonal within the PDX, the dissemination pattern was defined as monoclonal. Conversely, if any cluster defined as subclonal within the PDX sample was also present in the primary tumour, the divergence was classified as polyclonal.

If only a single PDX sample was considered for a patient, the patient level dissemination pattern matched the PDX level dissemination pattern. If multiple PDX models were sampled and the dissemination pattern of any individual PDX sample was defined as polyclonal, the patient level dissemination pattern was also defined as polyclonal. Conversely, if all PDX samples followed a monoclonal dissemination pattern all shared clusters between the primary tumour and each PDX were extracted. If all shared clusters overlapped across all PDX samples, the patient level dissemination pattern was classified as monoclonal, while if any PDX sample shared additional clusters with the primary tumour, the overall dissemination pattern was defined as polyclonal.

Furthermore, the origin of the seeding clusters was determined as monophyletic, if all clusters appear along a single branch, and polyphyletic if clusters were spread across multiple branches of the phylogenetic tree. Therefore, if a PDX sample was defined as monoclonal, the origin was necessarily monophyletic. For polyclonal PDX models, the clusters were mapped to branches of the evolutionary tree. If multiple branches were found, the origin was determined to be polyphyletic, while if only a single branch gave rise to all shared clusters the origin was defined as monophyletic.

For patient level definitions a similar approach was used. If any PDX was defined as polyphyletic, the overall origin was also defined as polyphyletic. Conversely, if all PDX samples were monophyletic in origin, all branches containing shared clusters were counted. If only a single such branch existed, the patient level origin was classified as monophyletic.

### Defining the seeding clones

The seeding clone is defined as the most-recent shared clone between the primary tumour and PDX model. Any cluster present in the primary tumour (defined as clonal or subclonal) and absent from the PDX sample was defined as primary specific, any cluster present solely in the PDX and absent from the primary tumour was defined as PDX specific, while all clusters present in both the primary tumour and PDX were defined as shared.

The shared clusters were mapped to the phylogenetic tree to determine the most recent shared cluster using a leaf-up approach. If the shared clusters could be mapped to a single branch of the phylogenetic tree, the clonality of the most recent shared cluster was determined in the PDX sample. If the most recent shared cluster was clonal in the PDX sample, this cluster was defined as the only seeding cluster for the PDX sample. On the other hand, if the most recent shared cluster was subclonal within the PDX, the parent cluster was also considered. This was done iteratively until the first shared cluster which was clonal in the PDX was found. Clusters along this path were defined as seeding if their phyloCCF value was greater than the phyloCCF of the child cluster.

If the shared clusters mapped to multiple branches of the phylogenetic tree, each branch was considered separately in the manner described above. If a parent cluster was shared between multiple branches, CCF values of both branches were added together and the iterative approach continued until the first cluster was found to be clonal in the PDX sample.

### Mutational distance

The mutational distance gives an approximation of mutational similarity between two regions, and also accounts for any large bottlenecks. Specifically, the distance will be large if few mutations are shared, or shared mutations occur at very different cellular frequencies; while the distance will be small if most mutations occur at similar frequencies across two regions.

Given two regions *i* and *j*, and *M* being the total number of mutations present in either one or the other region, excluding all truncal mutations; the mutation distance is calculated as:

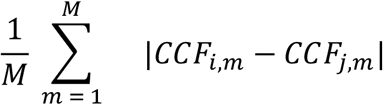

Where *CCF*_*i,m*_ and *CCF*_*j*.*m*_ are the CCF of mutation *m* in region *i* or *j*, respectively.

To calculate a distance for each region, the pairwise distance to each other region of interest is calculated and the average across all pairwise distances computed.

### Copy number distance

The copy number distance gives an approximation of similarity between two regions relating to relative gains and losses of segments. If gains/losses of segments relative to ploidy are consistent across two regions the copy number distance is small; whereas when they diverge, e.g. a loss in one region and neutral copy number state in the other, the distance increases.

### Statistical information

Statistical tests were performed in R (versions 3.6.3 & 4.1.1) or Prism 9.2.0. No statistical methods were used to predetermine sample size. Details of all statistical analyses are provided within figure legends. For all statistical tests, the number of data points included are plotted or annotated in the corresponding figure; and all statistical tests were two-sided unless otherwise specified.

## Data Availability

The whole exome sequencing data (from the TRACERx study and from PDX models derived from patients enrolled in TRACERx) used during this study have been deposited at the European Genome– phenome Archive (EGA), which is hosted by The European Bioinformatics Institute (EBI) and the Centre for Genomic Regulation (CRG) under the accession codes XXXXX; access is controlled by the TRACERx data access committee. Details on how to apply for access are available at the linked page. Biological materials, including PDX models generated within this study, will be made available to the community for academic non-commercial research purposes via standard MTA agreements upon publication.

## Code Availability

All code to reproduce figures can be found <u>here</u>. All code, unless already publicly available, will be made accessible upon publication. (https://zenodo.org/record/7434888?token=eyJhbGciOiJIUzUxMiIsImV4cCI6MTY4MzkzMjM5OSwiaWF0IjoxNjcwOTY4Nzg3fQ.eyJkYXRhIjp7InJlY2lkIjo3NDM0ODg4fSwiaWQiOjI4NDI3LCJybmQiOiIxMjQ1ZmI1NCJ9.EzByesRNyzrcLt13JI-6_3EKZ5v4u1O-q13d6q7Q75mK-0bIgQRHBAGBaFa9k-CpA72ghCV6hgwiYhhut_ifaw).

## ACKNOWLEDGEMENTS

The authors thank the staff of the Advanced Sequencing Facility at The Francis Crick Institute, as well as the members of the TRACERx consortium for their contributions to this study. TRACERx (Clinicaltrials.gov no: NCT01888601) is sponsored by University College London (UCL/12/0279) and was approved by an independent research ethics committee (REC 13/LO/1546). TRACERx is funded by Cancer Research UK (CRUK; C11496/A17786) and is coordinated by CRUK and the UCL Cancer Trials Centre. The authors thank Sharon Vanloo for administrative support. **R.E.H**. was supported by a Sir Henry Wellcome Postdoctoral Fellowship (Wellcome Trust; WT209199/Z/17) and received additional funding for this project from the CRUK Lung Cancer Centre of Excellence, the Roy Castle Lung Cancer Foundation and the James Tudor Foundation. R.E.H. is a National Institute for Health and Care Research (NIHR) Great Ormond Street Hospital (GOSH) Biomedical Research Centre (BRC) Collaborative Catalyst Fellow. The views expressed are those of the author(s) and not necessarily those of the NHS, the NIHR or the Department of Health. **T.K**. is supported by the Japan Society for the Promotion of Science (JSPS) overseas research fellowships program (202060447). **M.J.H**. is a CRUK Fellow and has received funding from CRUK, NIHR, the Rosetrees Trust, UKI NETs and the NIHR University College London Hospitals Biomedical Research Centre. **N.M**. is a Sir Henry Dale Fellow, jointly funded by the Wellcome Trust and the Royal Society (211179/Z/18/Z), and also receives funding from CRUK, the Rosetrees Trust, the NIHR BRC at University College London Hospitals and the CRUK University College London Experimental Cancer Medicine Centre. **C.S**. is a Royal Society Napier Research Professor (RSRP\R\210001). This work was supported by the Francis Crick Institute that receives its core funding from Cancer Research UK (CC2041), the UK Medical Research Council (CC2041), and the Wellcome Trust (CC2041). C.S. is funded by Cancer Research UK (TRACERx (C11496/A17786), PEACE (C416/A21999) and CRUK Cancer Immunotherapy Catalyst Network); Cancer Research UK Lung Cancer Centre of Excellence (C11496/A30025); the Rosetrees Trust, Butterfield and Stoneygate Trusts; NovoNordisk Foundation (ID16584); Royal Society Professorship Enhancement Award (RP/EA/180007); NIHR University College London Hospitals Biomedical Research Centre; the Cancer Research UK-University College London Centre; Experimental Cancer Medicine Centre; the Breast Cancer Research Foundation (US) BCRF-22-157; Cancer Research UK Early Detection and Diagnosis Primer Award (Grant EDDPMA-Nov21/100034); and The Mark Foundation for Cancer Research Aspire Award (Grant 21-029-ASP). This work was supported by a Stand Up To Cancer-LUNGevity-American Lung Association Lung Cancer Interception Dream Team Translational Research Grant (Grant Number: SU2C-AACR-DT23-17 to S.M. Dubinett and A.E. Spira). Stand Up To Cancer is a division of the Entertainment Industry Foundation. Research grants are administered by the American Association for Cancer Research, the Scientific Partner of SU2C. C.S. is in receipt of an ERC Advanced Grant (PROTEUS) from the European Research Council under the European Union’s Horizon 2020 research and innovation programme (grant agreement no. 835297). For the purpose of open access, the author has applied a CC BY public copyright licence to any Author Accepted Manuscript version arising from this submission.

