## Supplementary Figure for "Genomic evolution of non-small cell lung cancer patient-derived xenograft models"

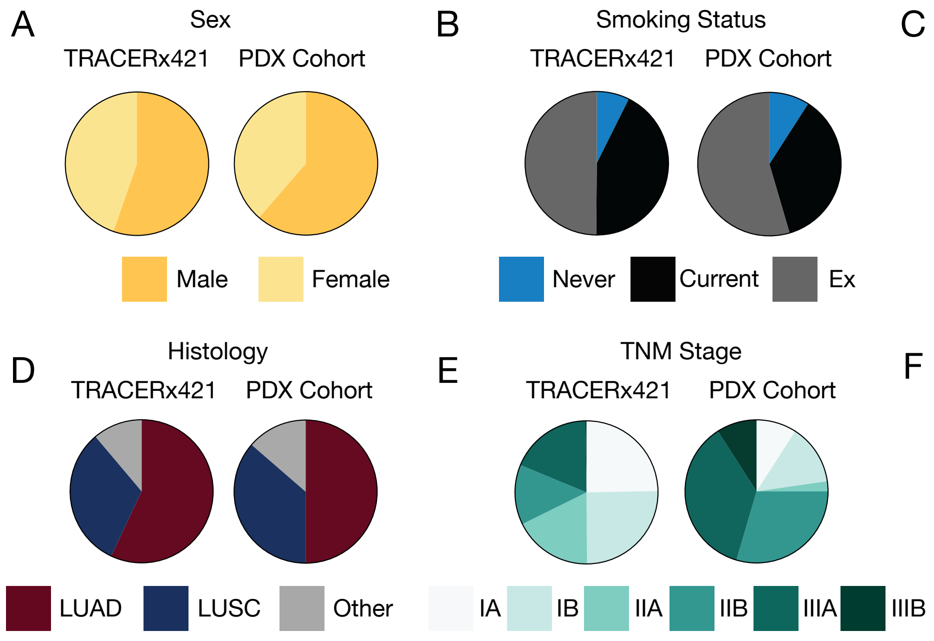


**Supplementary Figure 1: PDX cohort clinical characteristics.** Comparisons of the 44 patients from whom PDX models were attempted with the TRACERx421 cohort. A) sex, B) smoking status, C) adjuvant treatment status, D) Histology,  E) TNM stage, and F) Ethnicity.

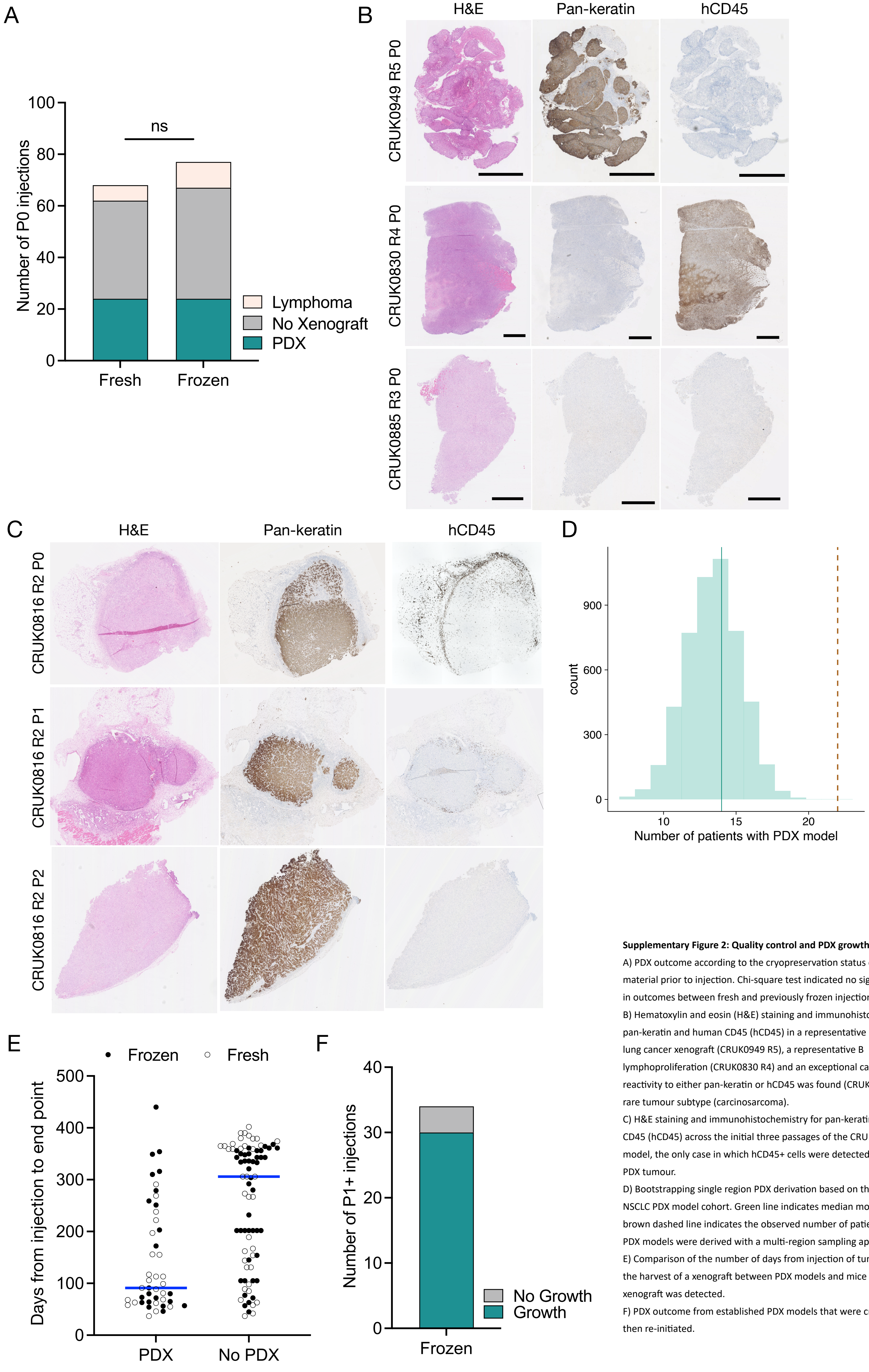


**Supplementary Figure 2: Quality control and PDX growth statistics.**

A) PDX outcome according to the cryopreservation status of primary tumour material prior to injection. Chi-square test indicated no significant difference in outcomes between fresh and previously frozen injections (p = 0.636).

B) Hematoxylin and eosin (H&E) staining and immunohistochemistry for pan-keratin and human CD45 (hCD45) in a representative non-small cell lung cancer xenograft (CRUK0949 R5), a representative B lymphoproliferation (CRUK0830 R4) and an exceptional case in with no reactivity to either pan-keratin or hCD45 was found (CRUK0885 R3), due to rare tumour subtype (carcinosarcoma).

C) H&E staining and immunohistochemistry for pan-keratin and human CD45 (hCD45) across the initial three passages of the CRUK0816 R2 PDX model, the only case in which hCD45+ cells were detected within the P0 PDX tumour.

D) Bootstrapping single region PDX derivation based on the described NSCLC PDX model cohort. Green line indicates median modelled solution, brown dashed line indicates the observed number of patients for whom PDX models were derived with a multi-region sampling approach.

E) Comparison of the number of days from injection of tumour material and the harvest of a xenograft between PDX models and mice in which no xenograft was detected.

F) PDX outcome from established PDX models that were cryopreserved and then re-initiated.

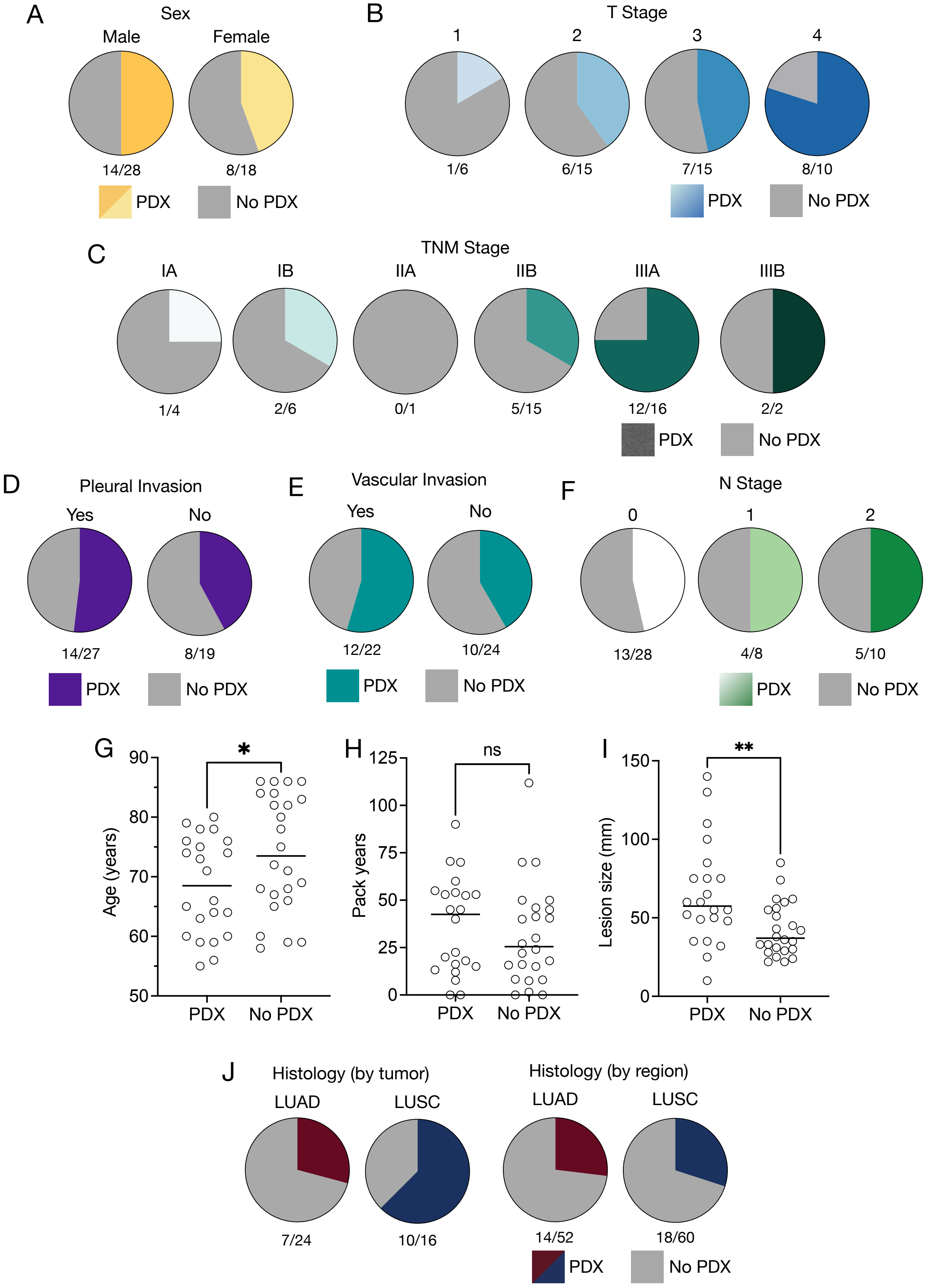


**Supplementary Figure 3: Patient characteristics as predictors of PDX engraftment.**

A) Comparison of patient sex distribution between tumors that generated no PDX models (grey = no PDX) and those that generated at least one PDX model (colour = PDX). B) Comparison of tumour stage (T stage) between those that generated no PDX models (grey = no PDX) and those that generated at least one PDX model (colour = PDX). C) Comparison of tumour stage (TNM stage, 8th edition) between tumours that generated no PDX models (grey = no PDX) and those that generated at least one PDX model (colour = PDX). D) Comparison of pleural invasion status between tumours that generated no PDX models (grey = no PDX) and those that generated at least one PDX model (colour = PDX). E) Comparison of vascular invasion status between tumours that generated no PDX models (grey = no PDX) and those that generated at least one PDX model (colour = PDX). F) Comparison of nodal stage (N stage) between tumours that generated no PDX models (grey = no PDX) and those that generated at least one PDX model (colour = PDX). G) Comparison of patient age between tumours that generated at least one PDX model and those that generated no PDX models (* = $p < 0.05$; two-tailed Mann-Whitney test, $p = 0.046$). H) Comparison of pack years of smoking between patients whose tumours generated at least one PDX model and those that generated no PDX models (ns = non-significant; two-tailed Mann-Whitney test, $p = 0.218$). I) Comparison of lesion size between tumours that generated at least one PDX model and those that generated no PDX models (** = $p < 0.01$; two-tailed Mann-Whitney test, $p = 0.004$). J) Comparison of histology between tumours (left) and regions (right) that generated no PDX models (grey = no PDX) and those that generated at least one PDX model (colour = PDX).

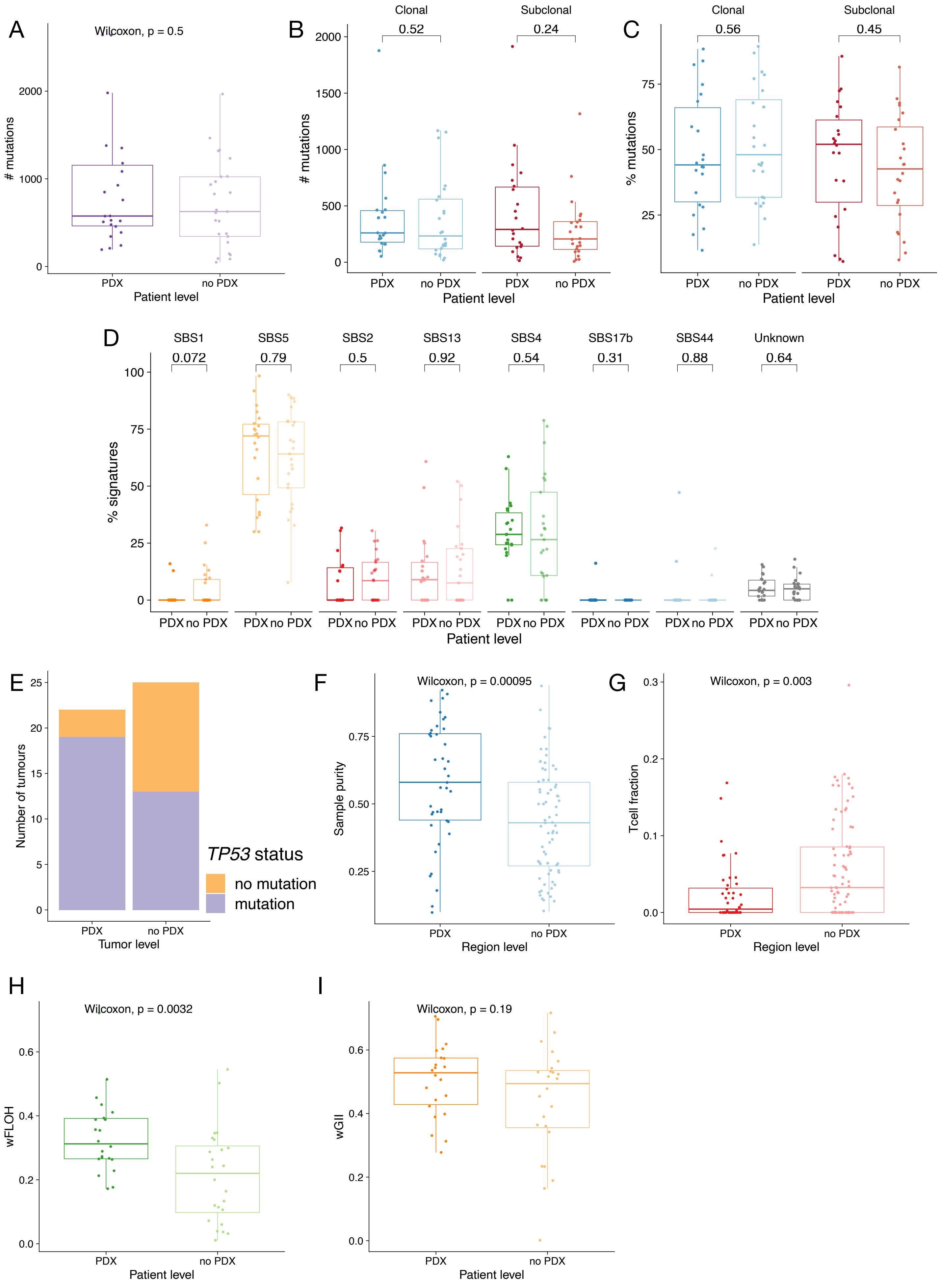
A
Wilcoxon, p = 0.5
### mutations
Patient level
B
Clonal
Subclonal
0.52
0.24
### mutations
Patient level
C
Clonal
Subclonal
0.56
0.45
% mutations
Patient level
D
SBS1
SBS5
SBS2
SBS13
SBS4
SBS17b
SBS44
Unknown
0.072
0.79
0.5
0.92
0.54
0.31
0.88
0.64
% signatures
Patient level
E
Number of tumours
Tumor level
TP53 status
no mutation
mutation
F
Wilcoxon, p = 0.00095
Sample purity
Region level
G
Wilcoxon, p = 0.003
Tcell fraction
Region level
H
Wilcoxon, p = 0.0032
wFLOH
Patient level
I
Wilcoxon, p = 0.19
wGII
Patient level
PDX
no PDX

**Supplementary Figure 4: Genomic characteristics as predictors of PDX engraftment.**

A) Comparison of total number of mutations between tumours that generated no PDX models and those that generated at least one PDX model (ns; two-tailed Wilcoxon rank sum test, p = 0.46).

B) Comparison of the number of truncal (ns; two-tailed Wilcoxon rank sum test, p = 0.53) and subclonal (ns; two-tailed Wilcoxon rank sum test, p = 0.18) mutations between tumours that generated no PDX models and those that generated at least one PDX model.

C) Comparison of the proportion of truncal (ns; two-tailed Wilcoxon rank sum test, p = 0.44) and subclonal (ns; two-tailed Wilcoxon rank sum test, p = 0.37) mutations between tumours that generated no PDX models and those that generated at least one PDX model.

D) Comparison of the mutational signature weights between tumours that generated no PDX models and those that generated at least one PDX model. No signatures were significant in a Wilcoxon rank sum test; p values shown.

E) Comparison of the *TP53* status of tumours that generated no PDX models and those that generated at least one PDX model (Fisher's exact test, p=0.015).

F) Comparison of the tumour purity of primary tumour regions that generated no PDX models and those that generated a PDX model (two-tailed Wilcoxon rank sum test, p = 0.0021).

G) Comparison of the adjusted T cell fraction of primary tumour regions that generated no PDX models and those that generated a PDX model (ns; two-tailed Wilcoxon rank sum test, p = 0.014).

H) Comparison of the weighted fraction of the genome subject to loss of heterozygosity (wFLOH) of primary tumours that generated no PDX models and those that generated a PDX model (two-tailed Wilcoxon rank sum test, p = 0.0032).

I) Comparison of the weighted genome instability index (wGII) score of primary tumours that generated no PDX models and those that generated a PDX model (two-tailed Wilcoxon rank sum test, p = 0.19).

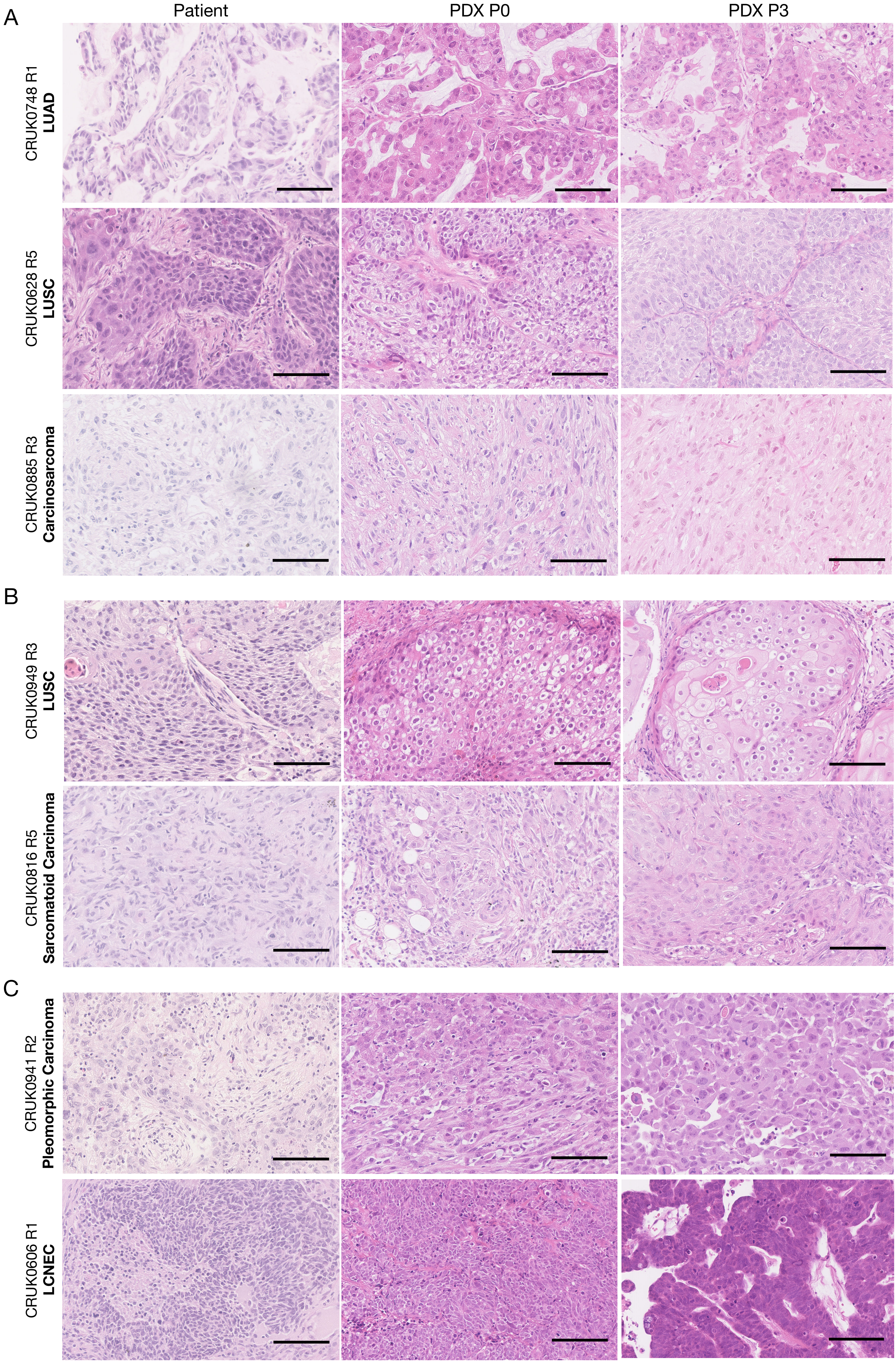

A
Patient
PDX P0
PDX P3
CRUK0748 R1
LUAD
CRUK0628 R5
LUSC
CRUK0885 R3
Carcinosarcoma
B
CRUK0949 R3
LUSC
CRUK0816 R5
Sarcomatoid Carcinoma
C
CRUK0941 R2
Pleomorphic Carcinoma
CRUK0606 R1
LCNEC

**Supplementary Figure 5: Histological comparison of patient tumour regions and matched PDX models.**

A) Examples of PDX models that retained histological similarity with the patient tumour at both early (P0) and late (P3) passage.

B) Examples of PDX models whose histological appearance diverged from the matched tumour region upon model establishment.

C) Examples of models whose histological appearance diverged from the matched tumour region at late passage (P3) having appeared similar at early passage (P0).

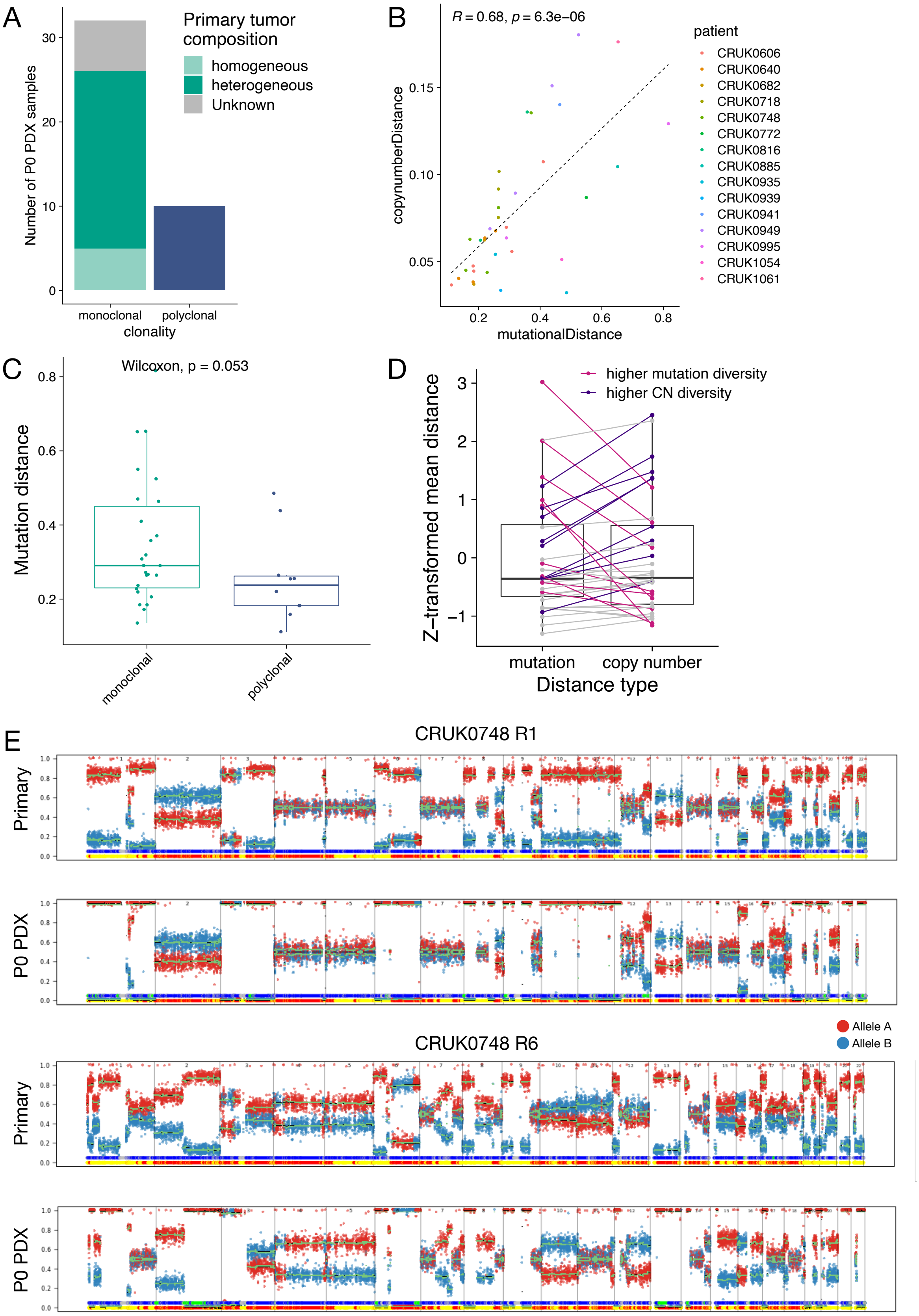


**Supplementary Figure 6: Related to Figure 3.**

A) Comparison of clonal composition of primary tumour regions giving rise to monoclonal PDX models.

B) Correlation between mutational distance and copy number distance. Points are coloured by patient ID. The black dashed line represents a linear model fitted to the points.

C) Comparison of mutational distance between region of origin and P0 for monoclonal versus polyclonal P0 PDX models.

D) Comparison of z-transformed mutation and copy number distance. Pink and purple lines represent the highest and lowest quartiles, respectively, of the difference between mutational and copy number diversity.

E) Example copy number profiles from CRUK0748 R1 and R6, which were identified as having higher mutational or copy number diversity, respectively. Allele A is coloured red, while allele B is coloured blue.

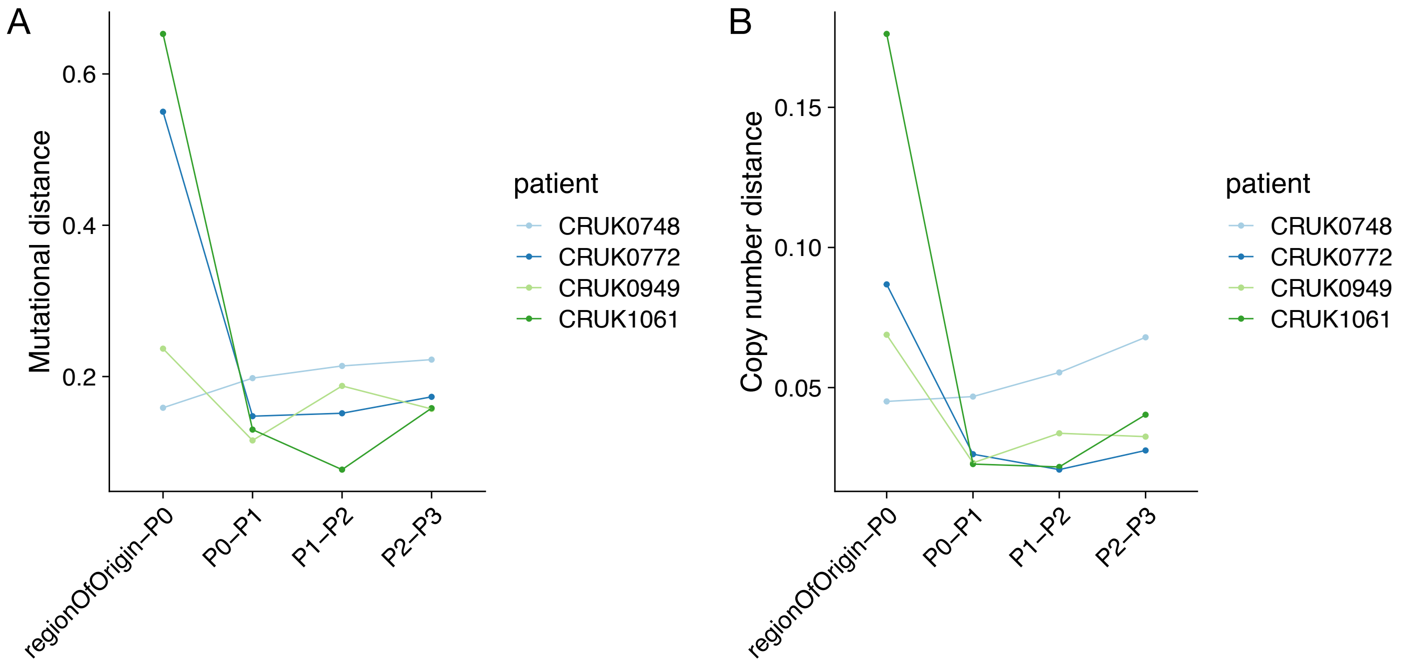


**Supplementary Figure 7: Related to Figure 5.**

A) Comparison of mutational distance over sequential passage in PDX samples.

B) Comparison of copy number distance over sequential passage in PDX samples.
